# cy3sabiork: A Cytoscape app for visualizing kinetic data from SABIO-RK

**DOI:** 10.1101/062091

**Authors:** Matthias König

## Abstract

Kinetic data of biochemical reactions are essential for the creation of kinetic models of biochemical networks. One of the main resources of such information is SABIO-RK, a curated database for kinetic data of biochemical reactions and their related information. Despite the importance for computational modelling there has been no simple solution to visualize the kinetic data from SABIO-RK.

In this work, I present cy3sabiork, an app for querying and visualization of kinetic data from SABIO-RK in Cytoscape. The kinetic information is accessible via a combination of graph structure and annotations of nodes, with provided information consisting of: (I) reaction details, enzyme and organism; (II) kinetic law, formula, parameters; (III) experimental conditions; (IV) publication; (V) additional annotations. cy3sabiork creates an intuitive visualization of kinetic entries in form of a species-reaction-kinetics graph, which reflects the reaction-centered approach of SABIO-RK. Kinetic entries can be imported in SBML format from either the SABIO-RK web interface or via web service queries. The app allows for easy comparison of kinetic data, visual inspection of the elements involved in the kinetic record and simple access to the annotation information of the kinetic record.

I applied cy3sabiork in the computational modelling of galactose metabolism in the Human liver.

## 1 Introduction

One of the main challenges for the modeling of biochemical systems is the availability of reliable information on the individual reaction steps and their kinetics from the literature. This information includes kinetic parameters with their rate equations as well as detailed descriptions of how these were determined {Wittig2014}.

SABIO-RK is a manually curated database for kinetic data storing comprehensive information about biochemical reactions and their kinetic properties, with data manually extracted from the literature and directly submitted from lab experiments {Wittig2012, Wittig2014}. Available information comprises kinetic parameters with their corresponding rate equations, kinetic law and parameter types and experimental conditions under which the kinetic data were determined. In addition, information about the biochemical reactions and pathways including their reaction participants, cellular location and the catalyzing enzyme are recorded {Wittig2012}.

The information in SABIO-RK is structured in datasets, so called database entries, which can be accessed either through the web-based user interface (http://sabio.h-its.org/) or via web services (http://sabiork.h-its.org/sabioRestWebServices). Both interfaces support the export of the data in the Systems Biology Markup Language (SBML), a free and open interchange format for computer models of biological processes {Hucka2003}.

Database entries are annotated with controlled ontologies and vocabularies based on Minimum Information Required for the Annotation of Models (MIRIAM {Laibe2007}), e.g. reaction participants (e.g. small chemical compounds and proteins), as well as kinetic rate laws, and parameters. The annotation information is encoded in the form of RDF-based MIRIAM annotations, and additional XML based SABIO-RK specific annotations, e.g. for experimental conditions. These annotations integrate the kinetic information with external resources like ChEBI {deMatos2010}, UniProtKB {UniProtConsortium2011}, Pubmed, or KEGG {Kanehisa2010}.

Despite the importance of kinetic information for computational modelling there has been no simple solution to visualize the database entries from SABIO-RK, and provide access to the network structure of the kinetic entries and the information encoded in the annotations.

In this work, we present cy3sabiork, an app for the visualization of kinetic data from SABIO-RK for Cytoscape, an open source software platform for network visualization {Shannon2003}. cy3sabiork creates an intuitive visualization of kinetic entries in the form of a species-reaction-kinetics graph extended with kinetic information, which reflects the reaction-centered approach of SABIO-RK. Hereby, the kinetic information is accessible via a combination of graph structure and annotations of nodes, with provided information consisting of: (I) reaction details, enzyme and organism; (II) kinetic law, formula, parameters; (III) experimental conditions; (IV) publication; (V) additional annotations. cy3sabiork allows for easy comparison of kinetic data, visual inspection of the elements involved in the kinetic record and simple access to the annotation information of the kinetic record.

## Methods

### Implementation

cy3sabiork was written in Java as an OSGi bundle for Cytoscape 3. The bundle activator adds the cy3sabiork Action to the Cytoscape icon bar, which provides access to the JavaFX based cy3sabiork dialog. The combination of Swing and JavaFX is implemented based on an JFXPanel with JavaFX GUI updates in Platform.runLater, Swing GUI updates via SwingUtilities.invokeLater. The GUI is a combination of web based components handled in a WebView and classical GUI components. SABIO-RK entries are retrieved via the web services using the RESTful API. SBML from the web service calls or the web interface export is imported using a Cytoscape Task created by the LoadNetworkFileTaskFactory. CyNetworks and CyNetworkViews for the imported kinetic entries are created by a CyNetworkReader registered for SBML files provided by cy3sbml {Koenig2012]}. During the app development the SBML CyNetworkReader was extended to support the SABIO-RK specific annotations and data. RDF based annotations are read with JSBML {Draeger2011} and hyperlinks to the respective resources are created by parsing the resources.

### Operation

An overview over the typical cy3sabiork workflow is depicted in **Figure 1**. The main steps of operation are (1) searching entries in SABIO-RK, (2) loading entries in cy3sabiork, and (3) visual exploration of results:

**Fig. 1.**
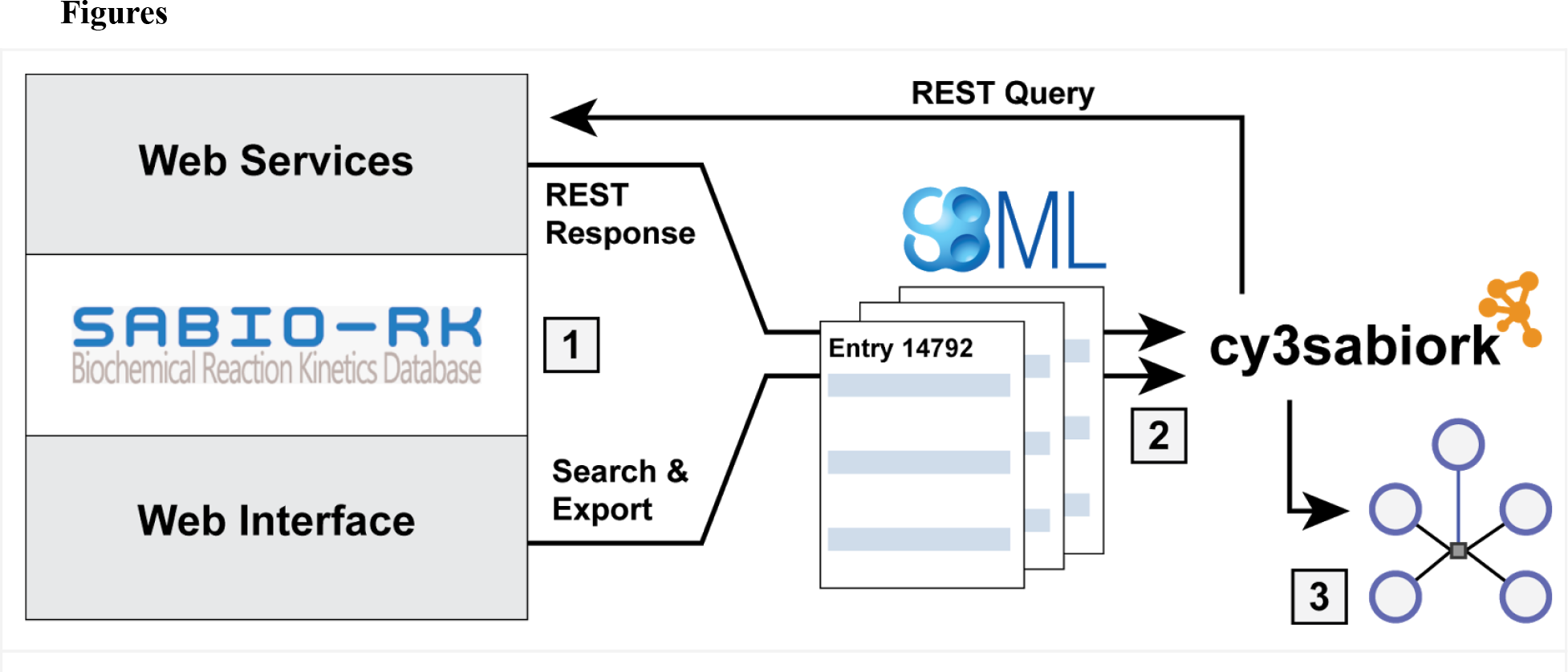
cy3sabiork workflow. The main steps of operation are (1) searching entries in SABIORK:Kinetic entries areretrieved from SABIORK via calls to the RESTful web services or via searching the web interface and exporting selected entries as SBML. (2) loading entries in cy3sabiork: The exchange format between SABIORKand cy3sabiork is SBML. (3)visual exploration of results. The kinetic graphs for the imported entries are generated providing simple access to kinetic information and annotations. An example query with resulting SABIORK information and subsequent visualization is shown in **Figure 2**.

#### *(1)* *Searching kinetic entries*

Kinetic entries can either be searched via the web services available from the cy3sabiork panel or directly in the SABIO-RK web interface available at http://sabio.h-its.org/.

The web-based user interface enable the user to search for reactions and their kinetics by specifying characteristics of the reactions. Beside a free text search, detailed queries can be executed in the advanced search. It offers the creation of complex queries by specifying reactions by their participants (substrates, products, inhibitors, activators etc.) or identifiers (KEGG or SABIO-RK reaction identifiers and KEGG, SABIO-RK, ChEBI or PubChem compound identifiers), pathways, enzymes, UniProt identifiers, organisms (NCBI taxonomy {Sayers2012}), tissues or cellular locations (BRENDA tissue ontology (BTO) {Gremse2011}), kinetic parameters, environmental conditions or literature sources {Wittig2012, Wittig2014}. After finalizing the search the selected kinetic entries are exported as SBML.

The web interface supports the same query options, which can be added in the REST query GUI, but allows a direct import of the SBML without the requirement for search in the web interface and export and subsequent import of the SBML.

Example queries are depicted in **Figure 2** and **Figure 4** with resulting SBML available as **Fupplementary File S1** and **F2**.

**Fig. 2.**
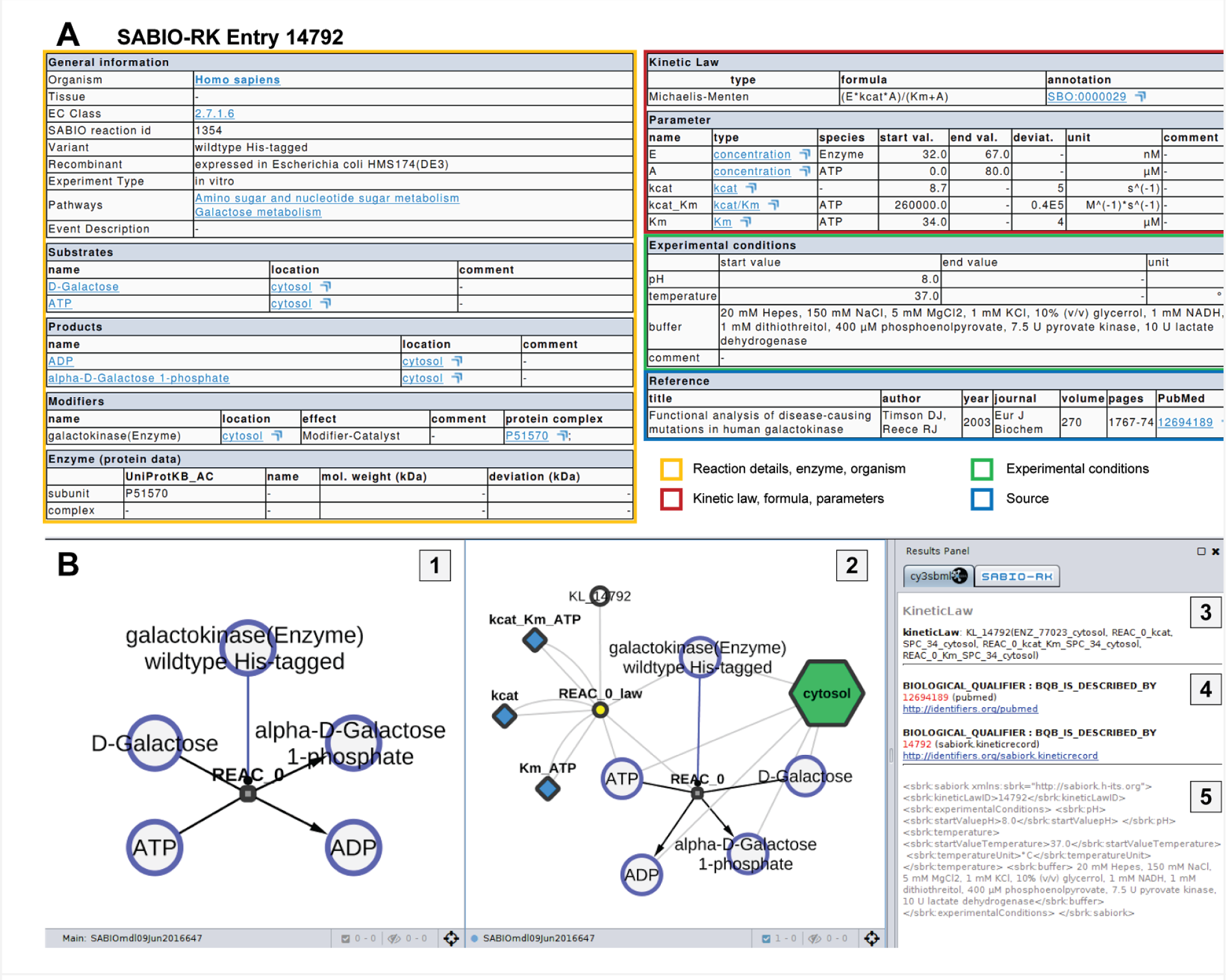
Overview of kinetic information and visualization for a single SABIORK entry. The kinetic entry 14792 for galactokinase (EC:2.7.1.6, UniProtKB:P51570) was retrieved via the web service query http://sabiork.hits.org/sabioRestWebServices/kineticLaws/14792 (status 10-06-2016, SBML of query provided as Supplementary File S1). (A) Overview of kinetic information for SABIORK entry (http://sabiork.hits.org/kineticLawEntry.jsp?viewData=true&kinlawid=14792) with color coding according to {Wittig2014}. (B) cy3sabiork information for entry 14792: [1] Resulting speciesreactionmodifier graph. The galacokinase enzyme catalyzes the conversion of DGalactose + ATP → α-D-Galactose 1phosphate + ADP (see also Substrates, Products and Modifiers in A). [2] kinetic graph with additional nodes for kinetic law, parameters and localization. [3] Selecting nodes in the graphs provides access to the annotation information and links to databases. In the example the kinetic law information is displayed in the Results Panel. [4] MIRIAM annotations with respective links to databases are available via the Results Panel. [5] Additional SABIORK annotations in XML for the experimental conditions are displayed in this section.

#### *(2)* *Loading kinetic entries*

The SBML exported from web interface searches is imported as in Cytoscape using cy3sbml {Koenig2012} (File → Import → Network → File). For queries to the web services the response SBML is imported automatically without the need for additional file operations.

#### *(3)* *Visual exploration*

The final step is the exploration of the kinetic entries in the species-reaction-modifier and the kinetic graph (e.g. in **Figure 2** and **Figure 4**). The information from a wide range of resources and databases is integrated with the graph visualization of the kinetic records accessible as hyperlinks from the cy3sbml panel. Examples are the access to the source publication on PubMed from which the kinetic information was retrieved, the UniProtKB protein for which the kinetic information was measured, KEGG and ChEBI information for species involved in the reaction, or links back to the SABIO-RK database entry and reaction.

## Use Cases

We applied cy3sabiork within the VLN and LiSyM projects for kinetic parameter search and model construction of a kinetic model of galactose metabolism of the human liver (https://github.com/matthiaskoenig/multiscale-galactose). A representative SABIO-RK query for galactokinase (EC:2.7.1.6, UniProtKB:P51570), the first step of hepatic galactose metabolization is depicted in **Figure 2** retrieving the SABIO-RK kinetic record 14792. A more complex query used for parameter search is depicted in **Figure 3** and **Figure 4** retrieving all kinetic records available for Human galactose metabolism.

During model building publications with kinetic information not yet available in SABIO-RK were included in the database by the SABIO-RK curation service.

**Fig. 3.**
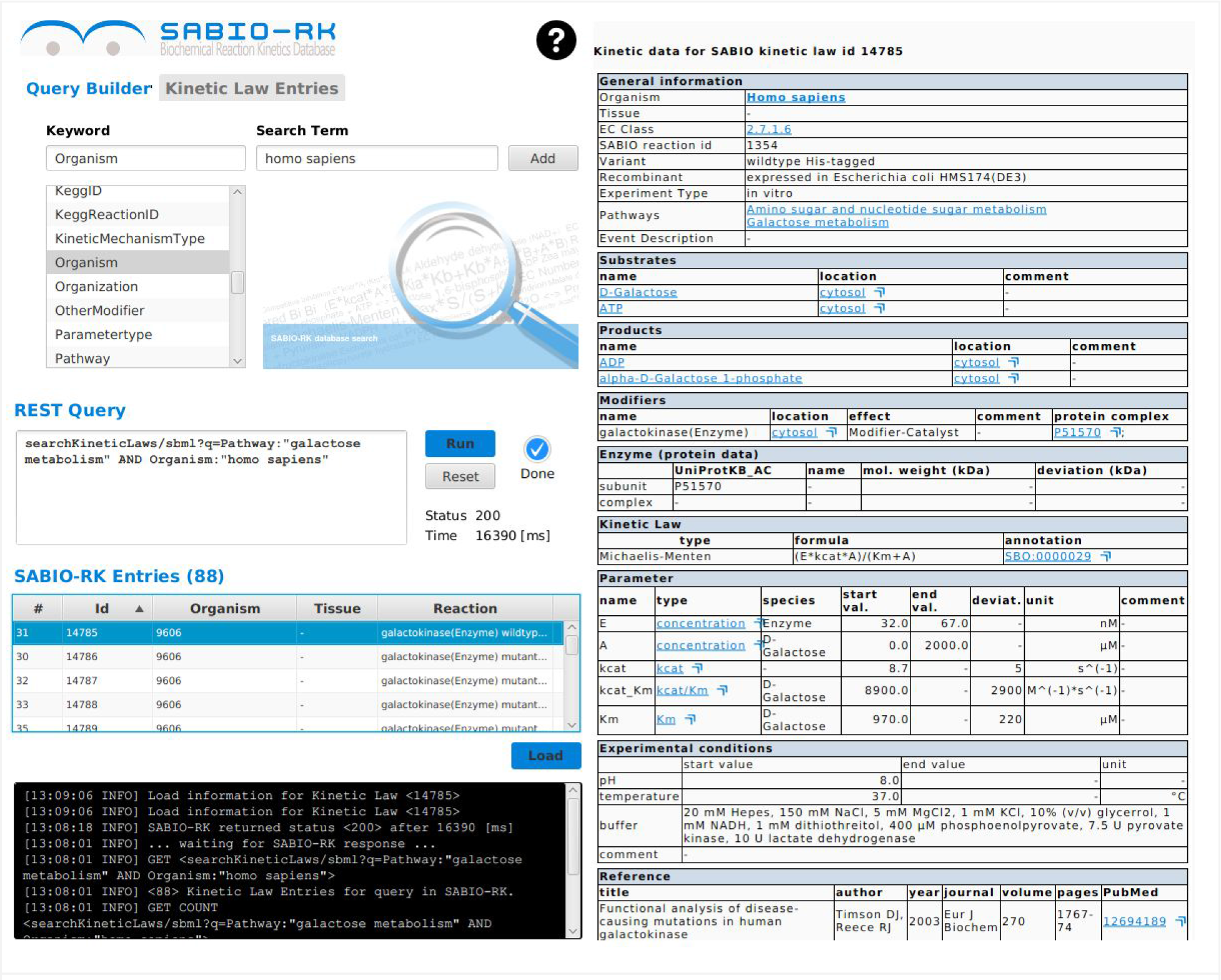
cy3sabiork GUI for web service queries. For Human galactose metabolism 88 entries are available (status 28-06-2016, SBML of query provided as **Supplementary File S2**). For the selected Entry 14785 detailed information is provided on the right side.

**Fig. 4.**
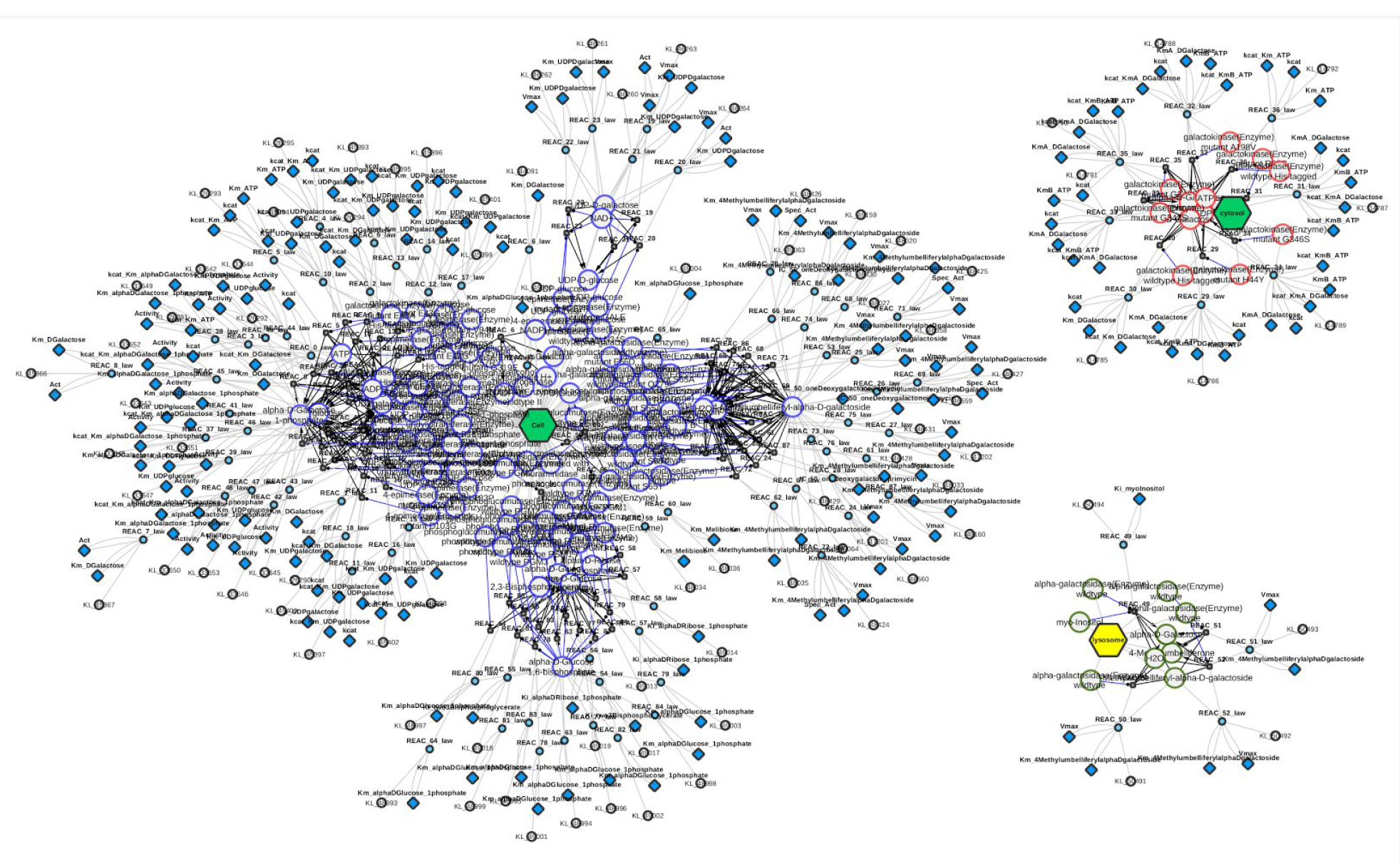
Graph of SABIORK kinetic information available for Human galactose metabolism consisting of 88 entries (status 28-06-2016, SBML of query provided as **Supplementary File S2**). The kinetic graph consists of three clusters, separated based on the reported localization of the catalyzing enzyme. The lysosomal entries are noncanonical reactions in the galactose metabolism.

## Conclusion

cy3sabiork is a Cytoscape app for visualizing kinetic data from SABIO-RK providing the means for visual analysis of kinetic entries from SABIO-RK within their reaction context. Herby, the integration of kinetic parameters with computational models is supported. The availability of direct links to annotated resources from within the network context of the kinetic records provides important information for the knowledge integration with computational models.

## Data and software availability

cy3sabiork is freely available from the Cytoscape App Store http://apps.cytoscape.org/apps/cy3sabiork. The code is open source under GNU General Public License, version 3 (GPL-3.0) available from the project homepage at https://github.com/matthiaskoenig/cy3sabiork.

## Author contributions

MK developed the app and wrote the manuscript. We thank the SABIO-RK team, the SBML community, and Cytoscape community for their support and help. A special thanks to the curation team of SABIO-RK including additional kinetic data in the database based on provided publications.

## Supplementary material

**Supplementary File S1**: SBML for query http://sabiork.h-its.org/sabioRestWebServices/kineticLaws/14792

**Supplementary File S2:** SBML for query http://sabiorkh-its.org/sabioRestWebServices/searchKineticLaws/sbml?a=Pathway:%22galactose%20metabolism%22%20AND%20Organism:%22homo%20sapiens%22

## References

De Matos, P., Alcántara, R., Dekker, A., Ennis, M., Hastings, J., Haug, K., et al. (2009). Chemical entities of biological interest: an update. Nucleic acids research, gkp886.

Dräger, Andreas et al. "JSBML: a flexible Java library for working with SBML." Bioinformatics 27.15 (2011): 2167–2168.

Gremse, Mario. et al. "The BRENDA Tissue Ontology (BTO): the first all-integrating ontology of all organisms for enzyme sources." Nucleic acids research 39.suppl 1 (2011): D507–D513.

Hucka, M., Finney, A., Sauro, H.M., Bolouri, H., Doyle, J.C., Kitano, H., et al. (2003). The systems biology markup language (SBML): a medium for representation and exchange of biochemical network models. Bioinformatics, 19(4), 524–531.

Kanehisa, M., Goto, S., Furumichi, M., Tanabe, M., & Hirakawa, M.(2010). KEGG for representation and analysis of molecular networks involving diseases and drugs. Nucleic acids research, J8(suppl 1), D355–D360.

König, M., Dräger, A., & Holzhütter, H.(2012). CySBML: a Cytoscape plugin for SBML. Bioinformatics, 25(18), 2402–2403.

Laibe, C., & Le Novère, N.(2007). MIRIAM Resources: tools to generate and resolve robust cross-references in Systems Biology. BMC Systems Biology, 1(1), 58.

Sayers, Eric W et al. "Database resources of the national center for biotechnology information." Nucleic acids research 37.Database issue (2009): D5.

Scheer, M., Grote, A., Chang, A., Schomburg, I., Munaretto, C., Rother, M., et al. (2010). BRENDA, the enzyme information system in 2011. Nucleic acids research, gkq1089.

Shannon, Pau. et al. "Cytoscape: a software environment for integrated models of biomolecular interaction networks." Genome research 13.11 (2003): 2498–2504.

UniProt Consortium (2011). Ongoing and future developments at the Universal Protein Resource. Nucleic acids research, 59(suppl 1), D214–D219.

Wittig, U., Kania, R., Golebiewski, M., Rey, M., Shi, L., Jong, L., et al. (2012). SABIO-RK—database for biochemical reaction kinetics. Nucleic acids research, 40(D1), D790–D796.

Wittig, U., Rey, M., Kania, R., Bittkowski, M., Shi, L., Golebiewski, M., et al. (2014). Challenges for an enzymatic reaction kinetics database. FEBS Journal, 281(2), 572–582.

